# Regimen comprising clarithromycin, clofazimine and bedaquiline is more efficacious than monotherapy in a mouse model of chronic *Mycobacterium avium* lung infection

**DOI:** 10.1101/2024.12.11.627976

**Authors:** Binayak Rimal, Ruth A. Howe, Chandra Panthi, Gyanu Lamichhane

**Author notes:** To whom correspondence should be addressed: Johns Hopkins University School of Medicine, 600 N Wolfe St, CMSC Room 3145, Baltimore, MD 21231.

## Abstract

*Mycobacterium avium*, a leading non-tuberculous mycobacterium (NTM) pathogen, causes chronic pulmonary infections, particularly in individuals with underlying lung conditions or immunosuppression. Current treatments involve prolonged multi-drug regimens with poor outcomes and significant side effects, highlighting the urgent need for improved therapies. Using a BALB/c mouse model of chronic *M. avium* pulmonary disease, we evaluated the efficacy of individual antibiotics— clarithromycin, clofazimine, and rifabutin—and combination regimens including clarithromycin+bedaquiline and clarithromycin+clofazimine+bedaquiline. Clarithromycin demonstrated potent bactericidal activity, reducing lung bacterial burden by 2.2 log_10_ CFU, while clofazimine transitioned from bacteriostatic to bactericidal, achieving a 1.7 log_10_ CFU reduction. Rifabutin was bacteriostatic against *M. avium* MAC 101 but ineffective against MAC 104. The triple-drug regimen of clarithromycin+clofazimine+bedaquiline was the most effective, achieving a 3.3 log_10_ CFU reduction in bacterial load, with 98% clearance within the first week and continued efficacy over eight weeks. Gross pathology confirmed these results, with granulomatous lesions observed only in untreated or rifabutin-treated mice. Combination therapy demonstrated enhanced efficacy compared to monotherapy. The findings underscore the potential of oral clarithromycin+clofazimine+bedaquiline or clarithromycin+clofazimine regimen as a promising therapeutic strategy for *M. avium* pulmonary disease.

## INTRODUCTION

*Mycobacterium avium* is a slow-growing, non-tuberculous mycobacterium (NTM) commonly found in water and soil (1). It is the most prevalent NTM pathogen in humans and a member of the *Mycobacterium avium* complex (MAC), which includes closely related species often indistinguishable using standard clinical microbiology staining techniques (2). *M. avium* primarily causes opportunistic lung infections, particularly in individuals with underlying lung comorbidities such as bronchiectasis, cystic fibrosis, or chronic obstructive pulmonary disease (COPD), as well as those with compromised immune systems (3). Most infections result from environmental exposure, and patients typically present with symptoms resembling bronchiectasis or tuberculosis (TB) (2). Relative to TB, caused by a related mycobacterium, treatment for *M. avium* pulmonary disease is challenging, with limited options and low cure rates (4).

The current standard of care involves multi-drug regimens comprising three or more antibiotics that inhibit essential functions in *M. avium* (5–8). Treatment typically lasts at least 18 months but may be further prolonged and is complicated by significant side effects, requiring frequent monitoring and adjustments. Despite these efforts, treatment outcomes remain poor. The increasing global prevalence of *M. avium* pulmonary disease underscores the urgent need for more effective and tolerable therapies. Only one drug, amikacin, has been approved for treating *M. avium* pulmonary disease based on a clinical trial (9). In contrast, TB drug development has progressed more rapidly, in part due to preclinical testing in animal models, particularly mouse models. These models have been critical in informing clinical trials for TB and other mycobacterial diseases (10).

To address this gap, Andrejak et al. developed a chronic *M. avium* pulmonary disease model using mice infected via aerosol exposure to mimic the natural infection route in humans (11). This model reproduces lung pathology in humans and has been validated with standard antibiotics such as clarithromycin, clofazimine, ethambutol, and rifampin, showing efficacy patterns consistent with human outcomes (11, 12). It has also been used to test experimental agents against *M. avium* (13). Using the BALB/c mouse model, we evaluated the efficacy of select antibiotics with *in vitro* activity against *M. avium* but uncertain effectiveness for lung disease. These included bedaquiline, clofazimine and rifabutin. Additionally, we assessed the efficacies of select drug combinations, as *M. avium* pulmonary disease typically requires regimens of three or more antibiotics. These included a two-drug combination of clarithromycin and bedaquiline and a three-drug combination of clarithromycin, bedaquiline, and clofazimine. The efficacy of the combination clarithromycin and clofazimine was not considered as it has been described using the same mouse model (12). Our study aims to identify more effective therapeutic options, addressing the critical need for improved treatments for *M. avium* pulmonary disease.

## RESULTS

### Monotherapy efficacies: Clarithromycin and clofazimine are bactericidal, rifabutin lacks efficacy

We assessed the efficacy of three antibiotics—100 mg/kg clarithromycin, 25 mg/kg clofazimine, and 20 mg/kg rifabutin—administered orally once daily to mice infected with MAC 101 (**Figure 1a**). At the time of infection, the mean lung bacterial load was 4.4 log_10_ CFU, which remained stable for four weeks before treatment began. In the control group treated with PBS (the solvent for the test antibiotics), the mean lung burden steadily increased over 12 weeks, resulting in a net increase of 0.95 log10 CFU, reflecting a steady, chronic infection.

**Figure 1:**
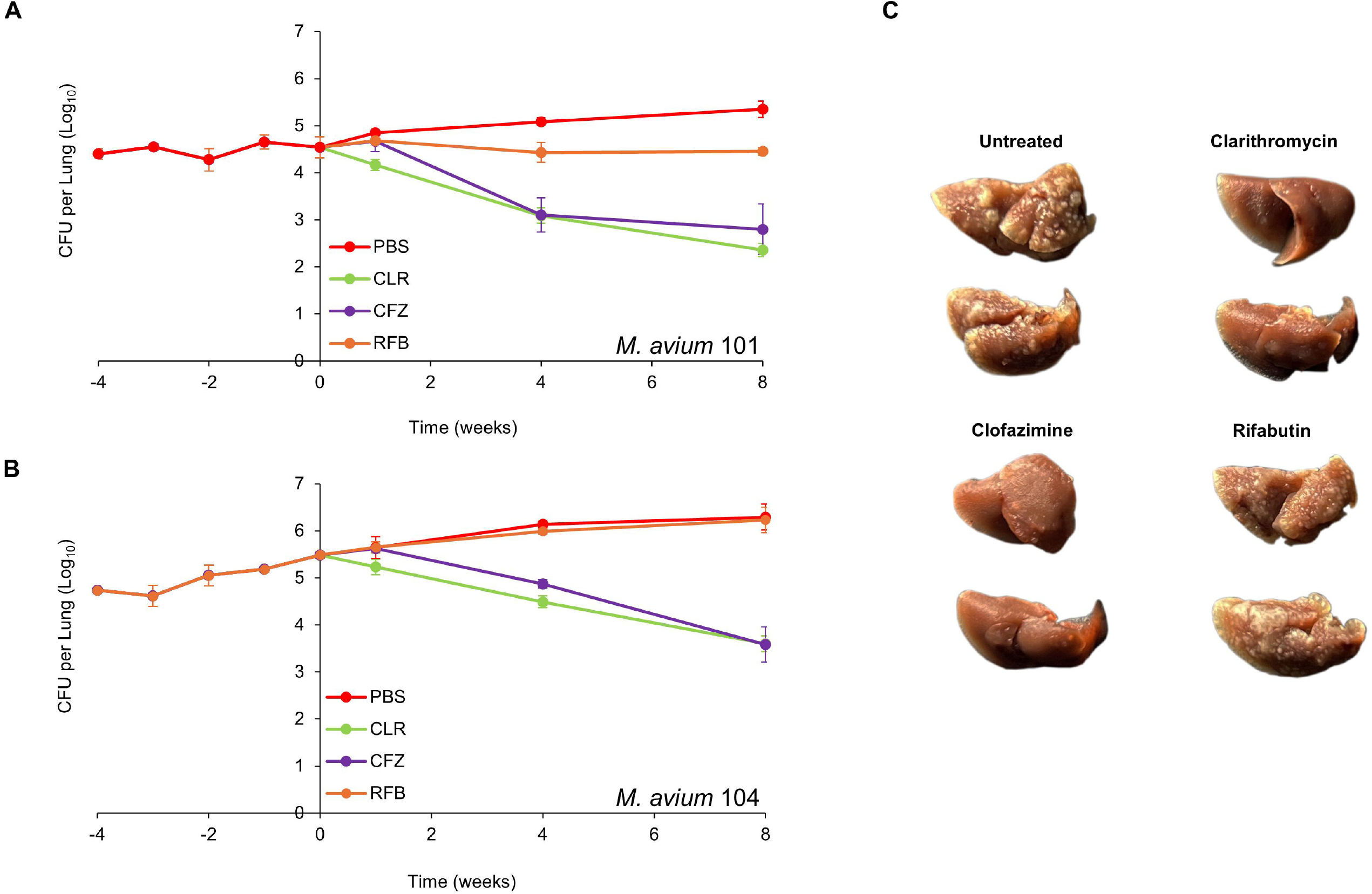
*M. avium* MAC 101 (A) and MAC 104 (B) burden in the lungs of BALB/c mice. Time point week -4 represents 24 h after infection with respective strain via the aerosol route. Time point week 0 represents conclusion of four weeks of infection and the day of antibiotic treatment initiation. Time points week-1, 4 and 8 represent the end of 1, 4 and 8 weeks of once daily oral administration of phosphate-buffered saline (PBS), 100 mg/kg clarithromycin (CLR), 25 mg/kg clofazimine (CFZ), and 20 mg/kg rifabutin (RFB). Mean CFU per lung and standard deviation are shown (n=5 per time point per group). (C) Gross pathology of the lungs of mice infected with MAC 104 from each treatment group, two mice per group at the conclusion of treatment (week 8) are shown.

In the rifabutin-treated group, the mean lung burden of MAC 101 remained stable throughout the treatment period, leading to a negligible net reduction of 0.09 log_10_ CFU after eight weeks. Thus, rifabutin displayed bacteriostatic activity against MAC 101. Statistical comparisons of the mean lung burden among treatment groups are provided in Table S1.

For clofazimine-treated mice, the lung burden remained unchanged after one week of treatment. However, at the end of four and eight weeks, net reductions in lung burden were 1.4 log_10_ CFU and 1.7 log_10_ CFU, respectively. This indicates that clofazimine initially exhibited bacteriostatic activity but became bactericidal with prolonged treatment. Clarithromycin demonstrated bactericidal activity from the start of treatment, achieving a net reduction of 2.2 log_10_ CFU by the end of the study. Among the three antibiotics, clarithromycin was the most effective against MAC 101.

A parallel experiment was conducted with mice infected with MAC 104 to validate the findings using an independent isolate (**Figure 1b**). At the time of infection, the mean lung burden was 4.7 log_10_ CFU, which increased by 1.5 log_10_ CFU over 12 weeks in the PBS-treated control group, consistent with a chronic infection. In rifabutin-treated mice, the lung burden followed a trajectory similar to the PBS group, indicating that rifabutin was ineffective against MAC 104. Clofazimine exhibited bacteriostatic activity during the first week of treatment but became bactericidal over time, producing a net reduction of 1.9 log_10_ CFU after eight weeks. Clarithromycin again demonstrated bactericidal activity throughout the treatment period, with a net reduction of 1.9 log_10_ CFU, matching the efficacy of clofazimine.

Gross pathological examination revealed granulomatous lesions in the lungs of mice treated with PBS or rifabutin, which were absent in mice treated with clarithromycin or clofazimine (**Figure 1c**). These pathological findings aligned with the microbiological results. In summary, rifabutin was bacteriostatic against MAC 101 but showed no activity against MAC 104. In contrast, clofazimine and clarithromycin were effective against both isolates, with clarithromycin being the most potent overall.

### Efficacy of regimen comprising clarithromycin, clofazimine and bedaquiline

The treatment of *M. avium* disease requires a multi-drug regimen to enhance efficacy and reduce the risk of selecting drug-resistant mutants (5–8). Consequently, neither clarithromycin nor clofazimine is used as monotherapy for this condition. However, given their strong anti-*M. avium* activity, we evaluated the efficacy of a regimen combining clarithromycin and clofazimine with a third agent, bedaquiline, in line with the current guideline recommendations to treat MAC lung infection with regimens comprising three or more agents. (5–8).

We tested a triple-drug regimen comprising 100 mg/kg clarithromycin, 25 mg/kg clofazimine, and 25 mg/kg bedaquiline against MAC 101 using the same protocol as described above (**Figure 2a**). In untreated mice, the lung burden of MAC 101 increased steadily, similar to the first study. The combination clarithromycin+clofazimine+bedaquiline demonstrated bactericidal activity throughout the treatment period, achieving a net 3.3 log_10_ CFU reduction in the lung burden of MAC 101. This represented a 98% reduction in bacterial load at the conclusion of the first week of treatment (**Figure 2b**). Of the remaining bacteria, 94% were cleared during the second to fourth weeks, and 54% of the survivors were eliminated in the final four weeks of treatment.

**Figure 2:**
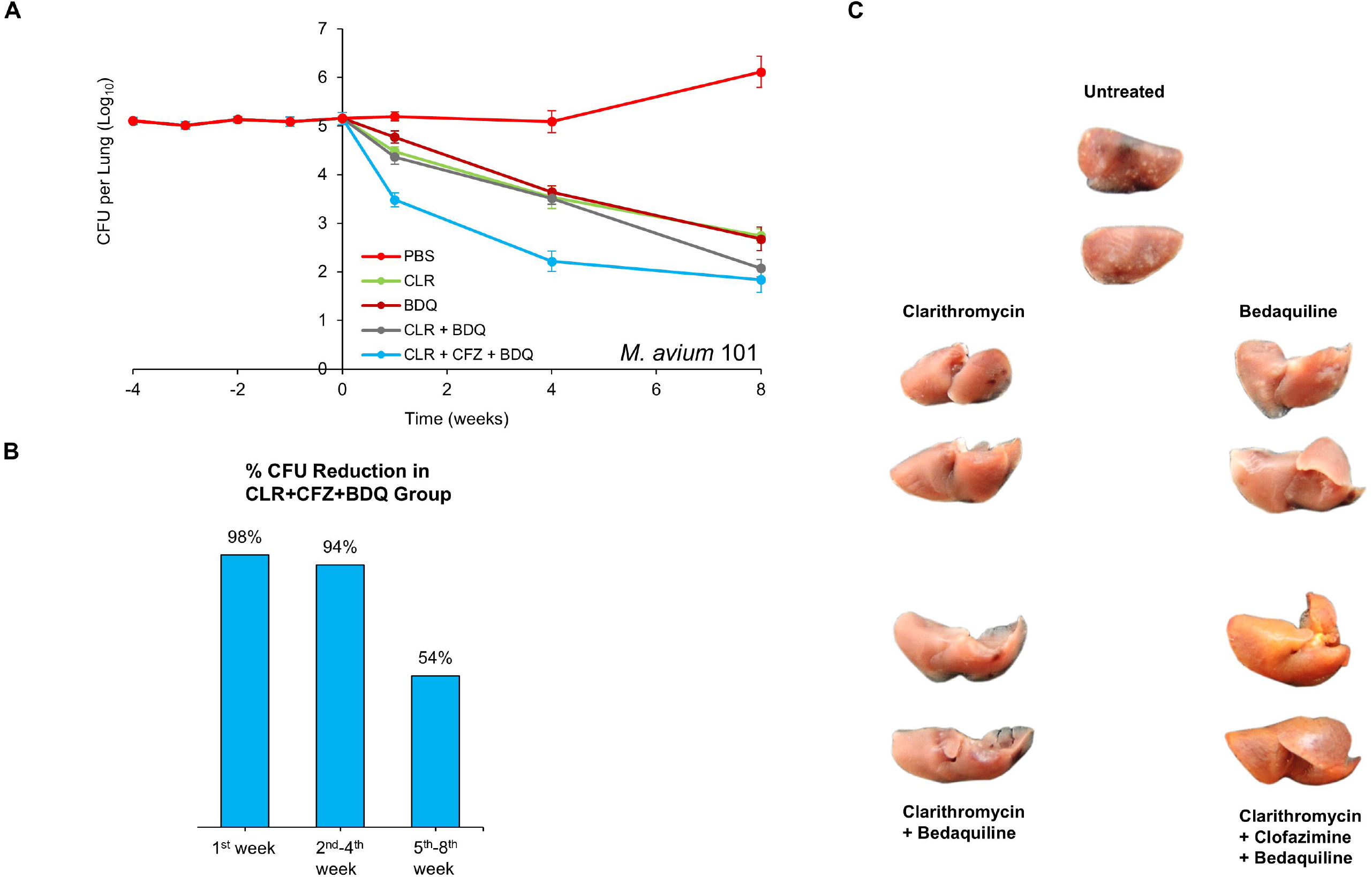
(A) *M. avium* MAC 101 burden in the lungs of BALB/c mice. Time point week -4 represents 24 h after infection via the aerosol route. Time point week 0 represents conclusion of four weeks of infection and the day of antibiotic treatment initiation. Time points week-1, 4 and 8 represent the end of 1, 4 and 8 weeks of once daily oral administration of phosphate-buffered saline (No treatment), 100 mg/kg clarithromycin (CLR), 25 mg/kg bedaquiline (BDQ) and 25 mg/kg clofazimine (CFZ). Mean CFU per lung and standard deviation are shown (n=5 per time point per group). (B) Percentage reductions in the mean MAC 101 burden in the lungs of mice treated with 100 mg/kg clarithromycin + 25 mg/kg clofazimine + 25 mg/kg bedaquiline during the first week, second-fourth week, and fifth-eighth week are shown. (C) Gross pathology of the lungs of mice infected with MAC 101 from each treatment group, two mice per group, at the conclusion of treatment (week 8) are shown.

Monotherapy with clarithromycin, clofazimine, or bedaquiline also reduced the MAC 101 lung burden, but at a slower rate compared to the triple-drug regimen (**Figure 2a and 1a**). The combination clarithromycin+bedaquiline was bactericidal throughout the treatment period, leading to a 3.1 log_10_ CFU reduction in lung MAC 101 burden. During the first four weeks of treatment, the addition of clofazimine significantly enhanced the potency of clarithromycin+bedaquiline, resulting in a greater reduction in lung burden. However, after eight weeks, both regimens produced statistically similar reductions in lung MAC 101 burden. This indicates that clofazimine primarily enhances the efficacy of clarithromycin+bedaquiline during the early stages of treatment, although paradoxically clofazimine monotherapy is bacteriostatic during this treatment stage.

Gross pathological examination at the end of the study revealed consolidated granulomas in the lungs of untreated mice (**Figure 2c**). These granulomas, a hallmark of *M. avium* lung disease in both mice (11) and humans (14), were absent in the lungs of mice treated with clarithromycin, bedaquiline, clarithromycin+bedaquiline, or clarithromycin+clofazimine+bedaquiline. Notably, the lungs of mice treated with the triple-drug regimen exhibited a reddish-yellow pigmentation, likely attributable to clofazimine, which is known to cause such pigmentation (15). Mice receiving antibiotics appeared healthy and showed no signs of sickness or lethargy throughout the study. In contrast, untreated mice became lethargic during the final stages of the study. Importantly, no deaths occurred in any of the treatment groups.

## DISCUSSION

Current treatment for MAC pulmonary infections is protracted and frequently complicated by the poor tolerability of complex regimens (4). Effective clinical decision-making, particularly when initiating treatment or modifying regimens to manage side effects, depends on a robust understanding of the bactericidal versus bacteriostatic efficacy of individual drugs and drug combinations. Unfortunately, such data has historically been more limited for MAC compared to TB (11, 12, 16–19). The kinetics of treatment response are critical clinical considerations, as therapy for chronic infections like MAC is often divided into distinct phases: a rapid-killing “induction” phase, followed by less intensive “consolidation” and “maintenance” phases. Each phase requires a dynamic balance between bactericidal efficacy, disease symptom management, mitigation of treatment side effects, and the logistical complexity of the regimen. Optimizing therapy to align with these shifting priorities at each phase has the potential to significantly enhance both patient experience and overall treatment outcomes.

Two distinct MAC isolates were included in this study to identify variations in drug efficacy, such as the differential activity of rifabutin, as well as instances where similar efficacies across isolates may allow for broader generalization of the findings to other strains. The dose and dosing frequency of bedaquiline, clarithromycin, clofazimine and rifabutin used in mice approximate their exposures in humans using approved doses. The treatment period was limited to eight weeks and was not designed to determine the duration required to achieve lung sterilization in mice. As such, the findings primarily offer valuable insights into the trajectory of early bactericidal activity associated with various regimens. This study focused on assessing drug efficacy against MAC isolates that are susceptible to bedaquiline, clarithromycin, clofazimine, and rifabutin. Furthermore, the main focus was to assess the efficacies of the dual combination clarithromycin+bedaquiline and the triple combination clarithromycin+clofazimine+bedaquiline that have not been evaluated before.

Consistent with clinical observations, rifabutin as a monotherapy displayed limited efficacy, showing only bacteriostatic activity at best against MAC 101 and no observable effect against MAC 104 (20, 21). On the other hand, Clarithromycin and clofazimine exhibited bactericidal activity against both MAC strains and were therefore tested in combination with bedaquiline. Again, consistent with prior observations for other mycobacteria, clofazimine as a monotherapy showed an initial bacteriostatic effect followed by delayed bactericidal activity (22, 23). Bedaquiline monotherapy closely paralleled the bactericidal trajectory of clarithromycin monotherapy by week four, although it showed comparably reduced bactericidal activity during the early stages of treatment. When clarithromycin was combined with bedaquiline, the regimen demonstrated early bactericidal activity similar to clarithromycin monotherapy, but with slightly more sustained bactericidal effects by week eight, indicating added benefit from the combination during later treatment stages.

The triple-drug combination of clofazimine, clarithromycin, and bedaquiline demonstrated a more rapid bactericidal effect against MAC 101 than was expected based on the effects of clofazimine monotherapy or the clarithromycin + bedaquiline dual therapy. This triple combination led to a greater than 1 log_10_ reduction in lung CFU burden at both weeks one and four, translating to a 98% reduction in organisms within the first week of treatment. This rapid early bactericidal activity contrasts sharply with the delayed bactericidal effect observed with clofazimine monotherapy against both MAC 101 and 104. However, this study did not evaluate clofazimine in two-drug combinations with either bedaquiline or clarithromycin, and as such, we cannot speculate whether these dual regimens might achieve comparable bactericidal timing to the three-drug combination. Previous study by Lanoix *et al*. suggested a synergistic relationship between clarithromycin and clofazimine based on their inclusion in more complex regimens alongside ethambutol and rifampin, though direct testing of clarithromycin and clofazimine as a standalone pair was not conducted (12). Future studies are needed to assess the efficacy of bedaquiline and clarithromycin in pairwise combinations with clofazimine.

The bactericidal trajectories observed in the two- and three-drug regimens in this study are both striking and clinically informative. At the conclusion of eight weeks of treatment, the clarithromycin+bedaquiline dual therapy achieved a level of bactericidal activity comparable to that of the clofazimine+clarithromycin+bedaquiline combination, although its early bactericidal effect was not as potent as that of the triple therapy. This finding suggests that clofazimine could be strategically added to or removed from a clarithromycin-based backbone, with or without bedaquiline, to tailor treatment across different phases. One potential approach would involve using the three-drug combination for its strong early bactericidal activity during the induction phase, then transitioning to clarithromycin+bedaquiline for the maintenance phase. Alternatively, if adverse side effects or drug-drug interactions pose significant concerns during the stabilization period, clofazimine could be introduced to a clarithromycin+bedaquiline regimen after stabilization. While beyond the scope of this study, future research on transitioning between such regimens at six to eight weeks of treatment could help further optimize bactericidal effects and inform clinical management strategies.

## MATERIALS AND METHODS

### Bacterial strains, growth media and growth conditions

*Mycobacterium avium* strain ATCC 700898, historically known as MAC 101, was purchased from American Type Culture Collection (Manassas, Virginia). *Mycobacterium avium* strain MAC 104 was a gift from Jacques Grosset laboratory, Johns Hopkins University, and used in the development of the mouse model of *M. avium* pulmonary disease (11). To infect mice, MAC 101 and MAC 104 were grown in Middlebrook 7H9 broth (Difco, catalog no. 271310) supplemented with 0.5% glycerol, 0.05% Tween-80 and 10% oleic acid-albumin-dextrose-catalase enrichment as described (24) in an orbital shaker at 220 RPM, 37 °C. MAC 101 and MAC 104 in the lungs of mice were grown by inoculating 10-fold serial dilutions of lung homogenates onto Middlebrook 7H11 selective agar (Difco, catalog no. 283810) supplemented with 0.5% glycerol, 0.05% Tween-80 and 10% oleic acid-albumin-dextrose-catalase enrichment (BD, catalog no. 212351), 50 μg/mL cycloheximide (Sigma-Aldrich, catalog no. C7698), and 50 μg/mL carbenicillin (Research Products International, catalog no. C46000).

### Antibiotics

All antibiotics preparations were made under sterile conditions. For clarithromycin (Sigma-Aldrich, catalog no. C9742), the amounts of the powder form necessary for each week of administration to mice were weighed into 50 ml polypropylene tubes prior to treatment initiation and stored at 4°C. At the beginning of each week, the weekly aliquot was retrieved, mixed with 0.05% agarose at 4°C to prepare a concentration of 10 mg/mL, and vortexed for 5 minutes. This preparation appears as white homogeneous suspension. The aliquot necessary for each day was transferred to 5 ml tubes and stored at 4°C until use. An 0.05% agarose solution was prepared by adding 50 mg Bacto agar (BD, catalog no. 214010) to 100 mL 1x phosphate buffered saline (PBS), pH 7.4 (Quality Biologicals, catalog no. 114-058-101), autoclaving for 10 min at 121°C and stored at 4°C until use.

For clofazimine (Sigma-Aldrich, catalog no. C8895), the weekly amount of powder was weighed into 50 mL polypropylene tubes and stored at 4°C. At the beginning of each week, the weekly aliquot was retrieved, mixed with 0.05% agarose at 4°C to prepare a concentration of 2.5 mg/mL, and vortexed for 5 minutes. This suspension was then sonicated at 50% power for 15 seconds per cycle, with 2-3 cycles, until a matte red, opaque, homogeneous colloidal suspension was achieved. Aliquots necessary for each day were transferred to 5 ml tubes and stored at 4°C until use.

For rifabutin (Sigma-Aldrich, catalog no. R3530), the amounts of powder necessary for each week were weighed into 50 ml polypropylene tubes prior to treatment initiation and stored at 4°C. At the beginning of each week, the weekly aliquot was retrieved, mixed with 0.05% agarose at 4°C to prepare a concentration of 2 mg/mL, and vortexed for 5 minutes. This preparation appears as dark red homogeneous suspension. The aliquot necessary for each day was transferred to 5 ml tubes and stored at 4°C until use.

For bedaquiline, powdered form bedaquiline fumarate (CAS no. 845533-86-0, Octagon Chemicals Ltd) was used. The amounts of the powder necessary for each week were weighed into a 100 ml borosilicate bottle, the precise volume of 20% 2-hydroxypropyl-β-cyclodextrin solution was added and dissolved by stirring with a magnetic stirrer for three hours at 4°C to prepare 2.5 mg/mL solution which appears transparent. Aliquots necessary for each day were transferred to 5 ml tubes and stored at 4°C until use. A 20% 2-hydroxypropyl-β-cyclodextrin (HPCD) (Sigma-Aldrich, catalog no. 332593) solution was prepared as described (25). Briefly, 20 g of HPCD powder was transferred to a 100-mL borosilicate bottle, and 75 mL of sterile deionized water was added and stirred with a magnetic stirrer until a clear solution was obtained (∼30 min). Approximately 1.5-mL of 1 N HCl was added to bring pH to 2.0, and the final volume was brought to 100 mL by adding sterile DI water. This solution was filtered through a 0.22-mm acetate cellulose filter and stored at 4°C until use.

### Infection and antibiotics efficacy assessment in mice

Three different cohorts of four-five weeks old female BALB/c mice were procured from the Charles River Laboratory (Wilmington, Massachusetts, USA) and housed in biosafety level 2 vivarium. Following arrival in our vivarium, mice were allowed to acclimatize for 7-10 days prior to initiating the studies. Mice were infected with MAC 101 or MAC 104 as described by Andrejak *et al* in a mouse model of *M. avium* lung infection (11). To infect mice, a fresh MAC 101 or MAC 104 culture at exponential phase, A_600nm_ of 1.00-1.60, was diluted in Middlebrook 7H9 broth to A_600nm_ of 1.0. 10 ml of this suspension was aerosolized with a nebulizer attached to Glas-Col Inhalation Exposure System A4212 (Glas-Col, Terre Haute, Indiana) into the chamber where all mice in an infection cohort were held. The infection sequence comprised of 15 minutes of pre-heat, 30 minutes of Mab suspension aerosolization into the chamber, 30 minutes of aerosol decay, and 15 minutes of surface decontamination with ultraviolet light. All mice in each study were infected simultaneously by natural breathing of the same *M. avium*-carrying aerosol for one hour.

To determine *M. avium* implantation in the lungs, five mice were sacrificed one day post infection (designated ‘week -4’), lungs were extracted aseptically, homogenized in 1xPBS with 2 mm glass beads by bead-beating for 30 seconds at 4,000 rounds-per-minute (Minilys, Bertin Instruments), 0.1 ml of appropriate 10-fold dilutions were inoculated onto selective Middlebrook 7H11 agar, incubated at 37 °C for 14 days and colony forming units were enumerated. Similarly, five mice were sacrificed at one-, two-, three- and four-weeks post infection (designated as weeks -3, -2, -1 and 0, respectively, in the figures) and lung *M. avium* burden was determined. Timepoint designated as ‘week 0’ represents the day antibiotics treatment was initiated and marks the conclusion of four weeks of infection. Lung *M. avium* burden was determined at the completion of one-, four- and eight-weeks of treatment (designated as ‘week+1, +4 and +8’, respectively) from five mice per treatment group, per timepoint.

Bedaquiline, clarithromycin, clofazimine and rifabutin were administered to deliver 25 mg/kg, 100 mg/kg, 25 mg/kg and 20 mg/kg of the antibiotics, respectively, per mouse, once daily, seven days a week for eight weeks. To achieve this, 0.2 ml bolus of 2.5 mg/ml bedaquiline, 10 mg/ml clarithromycin, 2.5 mg/ml clofazimine, and 2.0 mg/ml rifabutin preparations described were administered to each mouse by oral gavage using a 22-gauge curved gavage needle, with a 2-mm tip diameter (Gavageneedle.com; AFN2425C) fitted to a 1-mLslip-tip syringe (Becton & Dickinson, 309659).

### Ethics statement

Animal procedures described here were performed in adherence to the national guidelines and to the Johns Hopkins University Animal Care and Use committee approved protocol MO23M163.

### Lung Gross Pathology

In two efficacy assessment studies, one against MAC 101 and one against MAC 104, one half of the lungs from two mice from each treatment group at the final time point were allocated for lung gross pathology. Respective lungs were extracted, submerged in 5 ml 1x PBS for 48 hours and in 5 ml 10% buffered-formalin for 72 hours. The lungs were air dried and photographed.

### Data analysis

Raw lung CFU data were analyzed, and the mean ± standard deviation was calculated for each group at each timepoint. These results were graphed as dot plots. To assess the variance between treatment groups at each timepoint, a one-way ANOVA multi comparison was performed (**Table S1**), with significance determined at the 95% confidence level. A *p*-value of ≤ 0.05 was considered indicative of a non-random event, signifying significant differences in CFU burden between groups.

## AUTHOR CONTRIBUTIONS

BR: methodology, study design, investigation, data analysis and interpretation, manuscript preparation. RAH: data interpretation and manuscript preparation. CMP. Methodology and investigation. GL: study conception, study design, project administration, data interpretation, manuscript preparation, and funding acquisition.

